# Signed-XOR Error and Sparse Coding in a Dale-Compliant Substrate for Sequence Memorization

**DOI:** 10.64898/2026.06.24.734176

**Authors:** María Peña Fernández, Lara Lloret Iglesias, Jesús Marco de Lucas

## Abstract

How much machinery does a network need to memorize and recall discrete sequences when constrained to a biologically plausible substrate? We address this question using 50 short monophonic melodies in 4*/*4, used only as a controlled sequence-memory benchmark. Each beat is encoded with two clean one-hot populations – a 12-way pitch code and a separate 2-way {onset, sustain} code – and the decoder emits the same 14-dimensional code, so the autoregressive loop closes in a single neural format. The model obeys Dale’s law: latent units are excitatory or inhibitory, synaptic weights are non-negative, the decoder is implemented as explicit multi-contact bundles, and the encoder is a frozen sparse random projection wired at cortical (∼10%) density. On this substrate, a local ENGRAMMER signed-XOR read-out rule combined with a sparse *k*-winner-take-all code stores the training corpus exactly. With a modest latent expansion (*L* = 512), the model reaches 100% teacher-forced and autoregressive pitch accuracy, recognizes all training melodies, and separates all held-out melodies as novel with zero overlap. Ablations show that the signed error, sparse code, explicit E/I routing, and multi-contact synapses are the main load-bearing ingredients, whereas learning the encoder is strongly detrimental and dense input wiring does not help. Capacity sweeps show that Dale’s law mainly increases the capacity required for stable autoregressive recall: teacher-forced storage saturates between *L* = 128 and *L* = 256, while free-running recall becomes perfect by *L* = 512. A matched random corpus reaches the same final fidelity and is recalled at least as well at every capacity, indicating that musical structure does not improve recall on this benchmark and that final fidelity is set by capacity rather than by structure. The result is a Dale-compliant, gradient-free sparse associative memory rather than a general sequence learner.

## 1 Introduction

### 1.1 A controllable testbed for sequence memory

We want to know what a gradient-free network actually needs in order to store a set of sequences and recall them reliably *when it is held to a biologically constrained substrate*. To make that question crisp we need a task that is discrete, small enough that exact memorization is attainable, and unambiguous to score. A corpus of simple children’s melodies serves this purpose well, and we use it purely as such an example: the notes form a small vocabulary, each tune is short, and “did the model reproduce this tune” and “does the model treat this tune as familiar” are easy to quantify. We attach no significance to the musical content itself and draw no conclusions about how brains store songs or about long-term memory; the melodies are a convenient stand-in for any compact discrete sequence dataset.

### 1.2 From a static loss to a Dale-compliant substrate

The ENGRAMMER programme proposed the XOR motif as a candidate biological loss function: an inhibitory micro-circuit emits an error signal exactly when an internal prediction and an external signal disagree, and is silent when they match [1, 2]. As a local rule this trains a shallow autoencoder without backpropagation, and on binarized images the *signed* variant reconstructs inputs and organizes a separable latent space [2]. Applying the same signed rule to sequence memory, with a sparse *k*-WTA code and a frozen random encoder, is straightforward when the read-out weights are allowed to be *freely signed*; but a freely signed read-out is biologically unrealistic in two ways: real neurons are either excitatory or inhibitory and cannot flip the sign of their output (Dale’s law), and a single connection is not one signed scalar but a population of stochastic release sites. Here we therefore ask the sharper question: does the signed-XOR-plus-sparse-code recipe survive a substrate that obeys Dale’s law, uses explicit multi-contact synapses, and wires its encoder sparsely at cortical density, and if so, at what cost?

### 1.3 Approach and findings

We run the ablation constructively on the Dale’s-law substrate, turning each mechanism off in turn against a fixed reference, and only then vary capacity, connectivity, and the structure of the data itself. The findings are: (i) the signed error, the sparse *k*-WTA code, and the Dale population structure are each load-bearing; (ii) the encoder need not be learned, a frozen sparse random projection is best and a Hebbian encoder actively hurts; (iii) the input wiring can be cut to the ∼ 10% density of cortex at no cost, dense connectivity buying nothing, yielding a mushroom-body-like circuit; (iv) Dale’s law mainly raises the capacity required for stable free-running recall, not the attainable recall: teacher-forced storage saturates at 1.0 from *L* = 256, while autoregressive recall is the only metric that scales, reaching a perfect 1.0 by *L* = 512; and (v) against a matched random corpus, musical structure does not improve recall at any capacity (and is recalled at least as well by the random corpus at low capacity), so final fidelity is set by capacity rather than by structure. We then argue that the system is precisely a structured sparse associative memory, realized under Dale’s law.

## 2 Data: melodies in a 4/4 token format

The dataset is 62 monophonic children’s melodies in 4*/*4, partitioned into 50 training melodies and 12 held-out melodies that serve as a novelty probe. Each melody is segmented into bars of four beats. Every beat carries a pitch class (one of twelve) and an attack state (onset vs. sustain). We encode these as two separate one-hot populations rather than gluing a single attack bit onto the pitch code: a 12-way one-hot for the sounding pitch and a 2-way one-hot for {onset, sustain}. This keeps each field a clean one-hot, treats attack as its own attribute rather than a thirteenth pitch, and makes the code *symmetric* – every slot has exactly two active units, with sustain a positive state rather than the mere absence of an onset bit. The model reads *K* = 3 context bars and predicts the next bar; the flattened context is a 168-dimensional input vector (*K* × 4 = 12 slots, each a 14-dimensional code: pitch[12] + attack[2]), and the decoder emits the same 14-dimensional code, so the autoregressive loop closes in a single neural format (Sec. A compares this to a 13-dimensional attack-bit code). Across the 50 training melodies this gives 1,772 beat slots (1,445 onsets) and, with the *K* = 3 context window, 293 next-bar prediction examples.

Because the corpus is small and discrete, exact recall is attainable, so each mechanism’s effect can be read off the gap from 100% rather than from noisy generalization scores. We report pitch accuracy throughout; the attack channel is onset-dominated and reported only for completeness. (One held-out melody turns out to be impossible to store exactly, for a reason we return to in Section 4.7.)

## 3 Model and protocol

### 3.1 Substrate: a Dale’s-law encoder–decoder with a signed-XOR rule

The encoder maps the 168-d context to a latent **h** ∈ ℝ^*L*^ through a fixed random projection **W**_enc_; per-beat decoders read the next bar from **h** through **W**_dec_. Two biological constraints distinguish this substrate from the freely signed model. **Dale’s law**: each latent unit is excitatory or inhibitory in fixed proportion (80% E, 20% I); all weights are non-negative, and a unit’s influence on its targets carries the sign *s*_*j*_ ∈ {+1, − 1} of its *population*, so suppression must be routed through the inhibitory subpopulation rather than by a negative weight. **Quantal multi-synapse**: a decoder connection is an explicit bundle of up to ten release sites, each able to grow to unit strength; potentiation fills contacts and depression prunes them, so a many-contact connection transmits reliably while a one-weak-contact connection is labile. Learning is local and gradient-free: for each output unit a binary XOR mismatch *e*_*i*_ ∈ {0, 1} (1 when the prediction and target disagree) drives a Hebbian decoder update, with the update *direction* set jointly by the sign *c*_*i*_ ∈ {−1, +1} of the required correction and by the fixed Dale sign *d*_*j*_ ∈ {−1, +1} of the presynaptic latent unit *j* (Eq. 1):

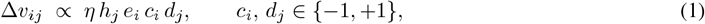

where *h*_*j*_ ≥ 0 is the presynaptic activity and *v*_*ij*_ is a contact bundle of **W**_dec_. Positive updates potentiate available contacts and negative updates prune labile contacts, with *v*_*ij*_ constrained to remain non-negative; suppression of an output is therefore implemented not by a negative weight but by potentiating the contacts from the inhibitory (*d*_*j*_ = − 1) population, which is how the Dale-compliant read-out routes negative corrections. A unit that should have fired but did not is potentiated and one that fired but should not is depressed, with the update routed onto contacts of the appropriate E or I population [2]. The encoder is *frozen* by default (a fixed random projection); we also test letting it learn through a fixed random feedback matrix (feedback alignment [3]).

### 3.2 Sparse code and sparse wiring

A hard *k*-winner-take-all nonlinearity (a fixed active count *k*, with *k/L* the code sparsity) keeps the *activity* sparse and decorrelated [4]. Separately, the input *wiring* is sparse: each latent unit draws a fixed random subset of the 168 inputs at a chosen connectivity density (default ∼ 10%, Section 4.5), so the encoder is a fixed *sparse* random projection. These are distinct: *k*-WTA sparsifies which units fire; sparse wiring sparsifies which inputs each unit can see at all. Together with the frozen random encoder this is the fly mushroom-body motif, sparse-random expansion read out by a local plastic rule [5, 6].

### 3.3 Regulatory mechanisms

Several mechanisms are switchable on top of the substrate, each a one-line ablation: synaptic *consolidation* by contact tagging (a repeatedly potentiated contact is locked and resists depression) [7, 8]; presynaptic-*LTD* pruning of the weakest labile contacts [9]; *divisive inhibition* normalizing the latent drive; and a *recognition* (song-ID) read-out head trained by the same signed rule. Recognition uses a vote over the per-window classifier head; novelty is the residual teacher-forced prediction error, thresholded against the known-song distribution. (A novelty-gated plasticity mechanism, in which already-known windows stop being rewritten [10, 11], is implemented in the code but is relevant only to incremental/continual learning and is not exercised in the static-corpus ablations reported here.)

### 3.4 Protocol

Each condition is a setting of the mechanism switches; the model code is unchanged across conditions. Capacity, connectivity, and ablation sweeps are trained for a fixed 120-epoch budget; the mechanism ablation is run at *L* = 256, *k* = 30 (the operating point where the full model is converged in teacher-forced storage but autoregressive recall still has headroom, so removals are discriminative). We report teacher-forced (TF) and autoregressive (AR) pitch accuracy, song recognition, and known–novel separation, as mean ± std over three seeds where noted (TF and recognition are effectively deterministic at saturation; AR is the high-variance measurement). A single fixed budget is justified by the training dynamics: at the reference configuration all metrics plateau well before 120 epochs (Fig. 3), so the budget is conservative headroom rather than a tuned quantity.

## 4 Results

### 4.1 Exact storage under Dale’s law

On the training corpus the Dale’s-law model, given a modest latent expansion (*L* = 512, *k* = 60), memorizes the corpus exactly: 100.0% teacher-forced and 100.0% autoregressive pitch accuracy (all 3 seeds), all 50 melodies recognized, and complete known–novel separation. The known-song residual is identically zero, while every one of the 12 held-out melodies has a clear positive residual, so the novelty threshold separates them with zero overlap. All 50 training melodies are reproduced perfectly under both teacher forcing and free running; the teacher-forced/autoregressive gap is exactly zero. This is the central result: the signed-XOR rule on a sparse code stores and recalls the corpus *flawlessly* even when the substrate is constrained to obey Dale’s law, uses explicit multi-contact synapses, and reads from a frozen encoder wired at 10% density. The remainder of this section establishes which ingredients are necessary (Sec. 4.2), how the result scales with capacity (Sec. 4.3) and connectivity (Sec. 4.5), and what musical structure does and does not contribute (Sec. 4.6).

### 4.2 What is load-bearing: the sign, the sparsity, and the E/I split

Turning each mechanism off in turn at *L* = 256 (Table 1, Fig. 1) – the operating point where the full model is converged (1.000 TF) but autoregressive recall has headroom (full reference AR 0.963 ± 0.027) – identifies four load-bearing ingredients and a clear set of inessential ones. Four changes collapse the model, taking down even teacher-forced storage. Removing the sparse *k***-WTA** code (0.571 ± 0.011 TF, 0.353 ± 0.019 AR) leaves a dense code from a random encoder too entangled for a local read-out to separate. **Learning the encoder** (Hebbian) is comparably destructive (0.391 ± 0.021 TF, 0.251 ± 0.012 AR, 0–3*/*12 novelty): error-driven learning belongs on the read-out, where the memory lives, not on the front end, which inverts the usual reading of feedback alignment as a stand-in for backpropagation [3, 12, 13]. **Collapsing the explicit E/I populations** (0.537 ± 0.044 TF, 0.310 ± 0.034 AR, novelty down to 3*/*12) removes the route by which a non-negative read-out can suppress an output, which is not in tension with a freely signed read-out – that implements negative corrections with negative weights, whereas here all weights are non-negative. And removing the **signed error / LTD channel** (LTP-only; 0.641 ± 0.047 TF, 0.365 ± 0.011 AR) shows that the directional ±1 error, not its magnitude, is what makes the local rule a usable descent direction.

**Table 1.** Leave-one-out ablation on the Dale’s-law substrate (50-melody training corpus, factored encoding, *L* = 256, *k* = 30, 120 epochs, mean ± std over 3 seeds). The full model is the frozen sparse-random encoder (10% wiring) with the signed-XOR multi-synapse decoder, Dale populations, *k*-WTA, consolidation, LTD, divisive inhibition, and the recognition head. Each row turns one mechanism off (or, last row, turns encoder learning on). Novelty is the count of the 12 held-out melodies flagged novel (range over seeds).

| Condition | TF pitch | AR pitch | Recog. | Novelty |
| --- | --- | --- | --- | --- |
| **full (reference)** | $1.000 \pm 0.000$ | $0.963 \pm 0.027$ | 1.000 | 12/12 |
| remove $k$-WTA sparsity | $0.571 \pm 0.011$ | $0.353 \pm 0.019$ | 0.880 | 6–11/12 |
| + encoder learning (Hebbian) | $0.391 \pm 0.021$ | $0.251 \pm 0.012$ | 0.800 | 0–3/12 |
| collapse E/I (positive-only single pop.) | $0.537 \pm 0.044$ | $0.310 \pm 0.034$ | 1.000 | 3–5/12 |
| remove the sign / LTD (LTP-only) | $0.641 \pm 0.047$ | $0.365 \pm 0.011$ | 1.000 | 9–12/12 |
| remove multi-synapse (scalar) | $0.998 \pm 0.002$ | $0.811 \pm 0.045$ | 1.000 | 12/12 |
| remove consolidation | $0.996 \pm 0.003$ | $0.905 \pm 0.046$ | 1.000 | 12/12 |
| remove divisive inhibition | $0.994 \pm 0.008$ | $0.898 \pm 0.064$ | 1.000 | 12/12 |
| remove sparse wiring (dense) | $1.000 \pm 0.000$ | $0.992 \pm 0.011$ | 1.000 | 12/12 |
| remove recognition head | $1.000 \pm 0.000$ | $0.981 \pm 0.006$ | — | 12/12 |

**Figure 1.**
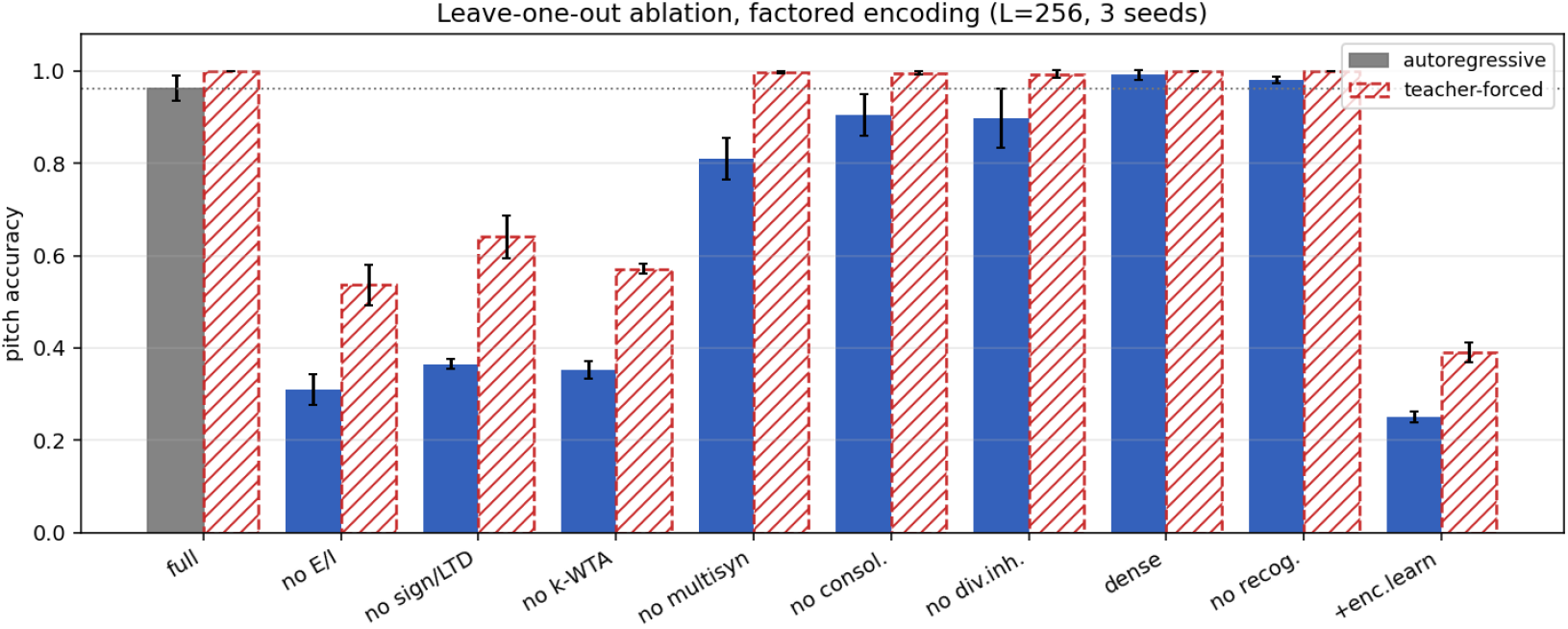
Leave-one-out ablation on the Dale’s-law substrate, factored encoding, at *L* = 256 (120 epochs, mean ± std over three seeds). Solid bars: autoregressive pitch accuracy; dashed hatched bars: teacher-forced; dotted line: the full-model AR reference. Removing *k*-WTA, letting the encoder learn, collapsing the E/I split, or removing the sign/LTD channel each collapses even teacher-forced storage; removing the multi-synapse representation materially lowers autoregressive recall; dense wiring and the recognition head are neutral for pitch.

The remaining mechanisms leave storage exact or near-exact and act only on autoregressive recall. The **quantal multi-synapse** representation materially lowers AR (to 0.811 ± 0.045 from the full 0.963) and divisive inhibition modestly so (0.898 ± 0.064); *consolidation* is marginal (0.905 ± 0.046). Two negative controls close the picture: **dense input wiring buys nothing** – the fully connected encoder is statistically indistinguishable from the 10% default (0.992 ± 0.011 vs 0.963 ± 0.027 AR) – and the recognition head is neutral for pitch (0.981 ± 0.006).

### 4.3 Capacity scaling: Dale’s law raises the cost of free-running recall

Sweeping the latent size *L* ∈ {64, 128, 256, 512, 1024}, with the active count *k* increased approximately in proportion (so that the code stays in a fixed sparse-coding regime, *k/L* ≈ 0.12, up to *k* = 60, beyond which *k* is held at 60; Table 2, Fig. 2), cleanly separates two regimes. A *storage floor* sits between *L* = 128 and *L* = 256: at *L* = 128 the model cannot yet hold all 293 associations (0.750 TF), and at *L* = 64 the code is too small to store the corpus (0.476 TF) or to reliably reject novelty (only 4–6 of 12 held-out melodies flagged), although recognition remains perfect throughout; by *L* = 256 teacher-forced storage is *exact* (1.000) and stays there. Above the floor, teacher-forced storage, recognition, and novelty are all saturated, and *autoregressive recall is the only metric that scales with L*: it climbs from 0.963 ± 0.027 at *L* = 256 to a perfect 1.000 ± 0.000 at *L* = 512 (all 3 seeds), and remains perfect at *L* = 1024. The interpretation is that Dale’s law mainly raises the *capacity required* for stable free-running recall rather than lowering the attainable recall: a positive-only excitatory/inhibitory read-out, which must route every suppression through the inhibitory subpopulation, needs a modest latent expansion to widen its competitor margins to the point of perfect free-running recall, and given those units it gets there exactly. The storage floor itself lies between *L* = 128 and *L* = 256 *within* an approximately fixed sparse-code regime (*k/L* ≈ 0.12): since *L* = 128 is already at that sparsity yet does not store the corpus, the limiting factor is not sparsity alone but the joint geometry of code dimensionality and active-set size. The threshold is thus representational (it is not crossed by simply lowering *k/L*), and it is not a matter of raw parameter count either.

**Table 2.**
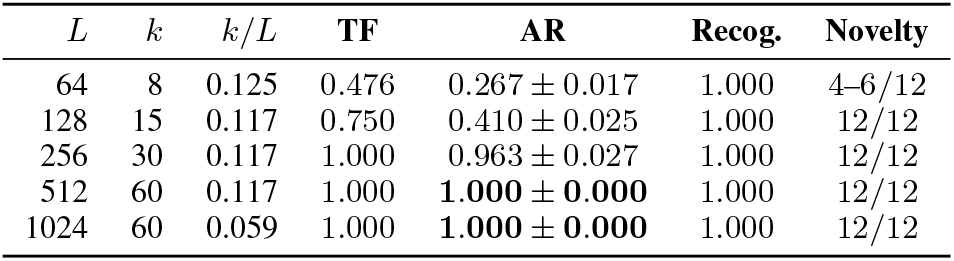
Capacity sweep on the training corpus, factored encoding (120 epochs, mean ± std over 3 seeds). Teacher-forced storage, recognition, and novelty saturate above the storage floor; autoregressive recall is the only metric that scales, reaching a perfect 1.000 by *L* = 512.

| $L$ | $k$ | $k/L$ | TF | AR | Recog. | Novelty |
| --- | --- | --- | --- | --- | --- | --- |
| 64 | 8 | 0.125 | 0.476 | $0.267 \pm 0.017$ | 1.000 | 4–6/12 |
| 128 | 15 | 0.117 | 0.750 | $0.410 \pm 0.025$ | 1.000 | 12/12 |
| 256 | 30 | 0.117 | 1.000 | $0.963 \pm 0.027$ | 1.000 | 12/12 |
| 512 | 60 | 0.117 | 1.000 | **$1.000 \pm 0.000$** | 1.000 | 12/12 |
| 1024 | 60 | 0.059 | 1.000 | **$1.000 \pm 0.000$** | 1.000 | 12/12 |

**Figure 2.**
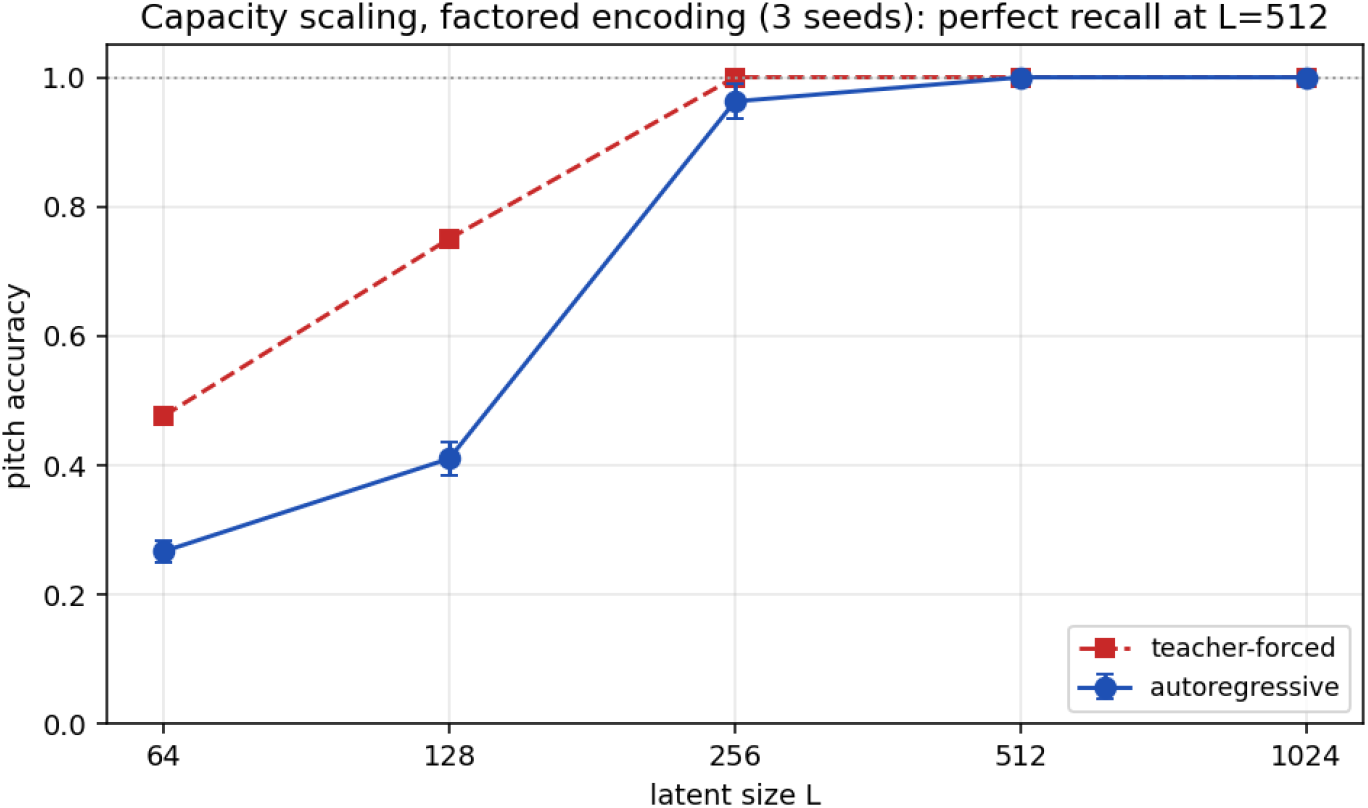
Capacity scaling on the Dale’s-law substrate (training corpus). Teacher-forced storage (open markers) is exact from *L* = 256; autoregressive recall (filled) is the only metric that scales, reaching a perfect 1.0 at *L* = 512. Below *L* = 256 the code is too small to store the corpus (storage floor); the floor lies within an approximately fixed sparse-code regime (*k/L* ≈ 0.12), so it is not crossed by sparsity alone.

### 4.4 Training dynamics: how many epochs are needed

The 120-epoch budget is deliberately conservative; tracking the reference configuration (*L* = 512, *k* = 60, factored) over training shows that far fewer epochs suffice (Fig. 3, mean over 3 seeds). Teacher-forced storage is the first to converge – it reaches ∼ 0.96 by epoch 30 and is essentially exact (≥ 0.995) by epoch 40 – while autoregressive recall, the slower and more demanding metric, lags behind: it passes 0.97 by epoch 40, 0.99 by epoch 52, and saturates at a perfect 1.000 by epoch ∼ 60. Beyond 60 epochs nothing changes. The ordering mirrors the capacity result: storage settles quickly, and the extra epochs (like the extra latent units) go into widening the basins until every free-running roll-out stays on the stored manifold. In practice ∼ 60 epochs reproduce the headline result exactly and ∼ 30 epochs already give near-perfect teacher-forced storage; we report the 120-epoch budget throughout only so that the slowest configurations in the sweeps are unambiguously converged.

**Figure 3.**
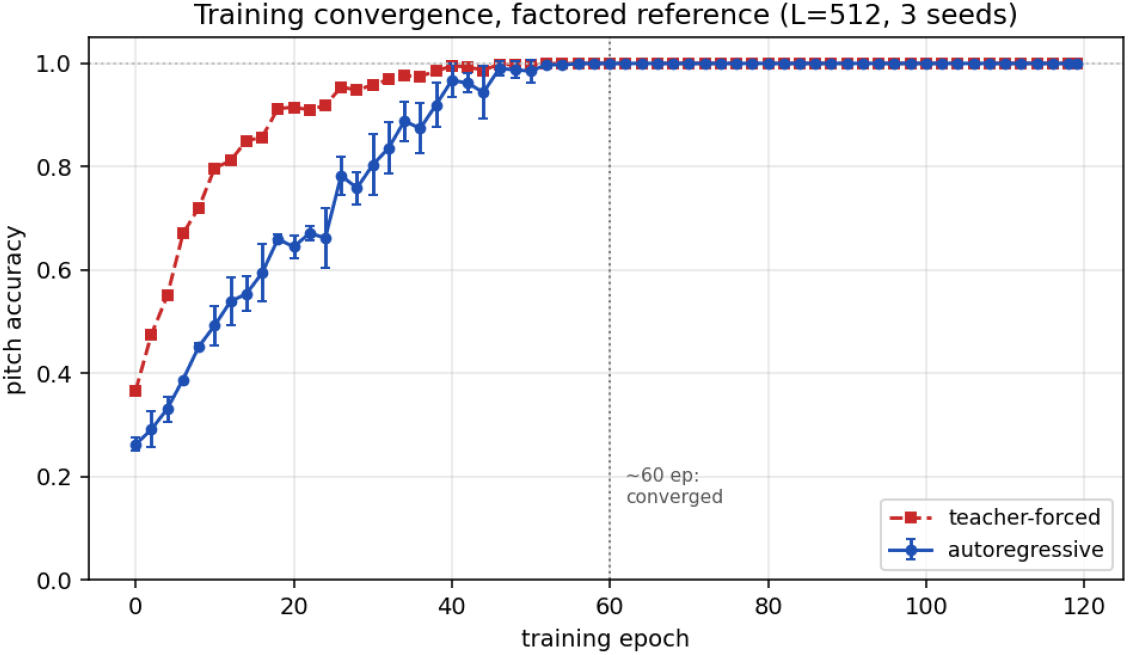
Training convergence at the reference configuration (*L* = 512, *k* = 60, factored encoding; mean ± std over 3 seeds). Teacher-forced storage (dashed) is essentially exact by epoch ∼ 40; autoregressive recall (solid) saturates at a perfect 1.000 by epoch ∼ 60, after which training has no further effect. The 120-epoch budget used elsewhere is roughly 2× this and serves only as conservative headroom.

### 4.5 Sparse input wiring at cortical density

Cortical neurons contact only a small fraction of possible partners (local excitatory connection probabilities on the order of 10% [14]), and the fly mushroom body wires each Kenyon cell to only ∼ 6–7 of ∼ 50 inputs [5, 6]. We therefore sweep the encoder’s wiring density at *L* = 256 (Table 3, Fig. 4; mean ± std over 3 seeds). Teacher-forced storage stays exact (1.000) from dense down to the ∼ 10% cortical default, and autoregressive recall is robust across the same range: it holds at 0.96–1.00 from dense to 10%, with these densities overlapping within seed variance, and only at 5% does it fall (0.823 ± 0.103, with storage itself beginning to drop, 0.966 TF). We do not read the small ordering among the higher densities as meaningful: across seeds, dense (0.992 ± 0.012) and the 10% default (0.963 ± 0.027) are statistically indistinguishable. The robust conclusion is simply that cutting the encoder’s wiring to cortical density carries no measurable cost. The working model is therefore a fixed sparse random expansion at cortical density read out by a local signed rule under Dale’s law, i.e. a mushroom-body-like associative memory, and dense connectivity buys nothing over it.

**Table 3.** Input-connectivity sweep at *L* = 256, *k* = 30, factored encoding (120 epochs, mean ± std over 3 seeds). Fixed random wiring at the given density; *k*-WTA activity sparsity is unchanged. Recognition 50/50 and novelty 12/12 throughout.

| Wiring density | TF pitch | AR pitch |
| --- | --- | --- |
| 100% (dense) | 1.000 | $0.992 \pm 0.012$ |
| 25% | 1.000 | $1.000 \pm 0.000$ |
| 15% | 1.000 | $0.996 \pm 0.006$ |
| 10% (default) | 1.000 | $0.963 \pm 0.027$ |
| 5% | 0.966 | $0.823 \pm 0.103$ |

**Figure 4.**
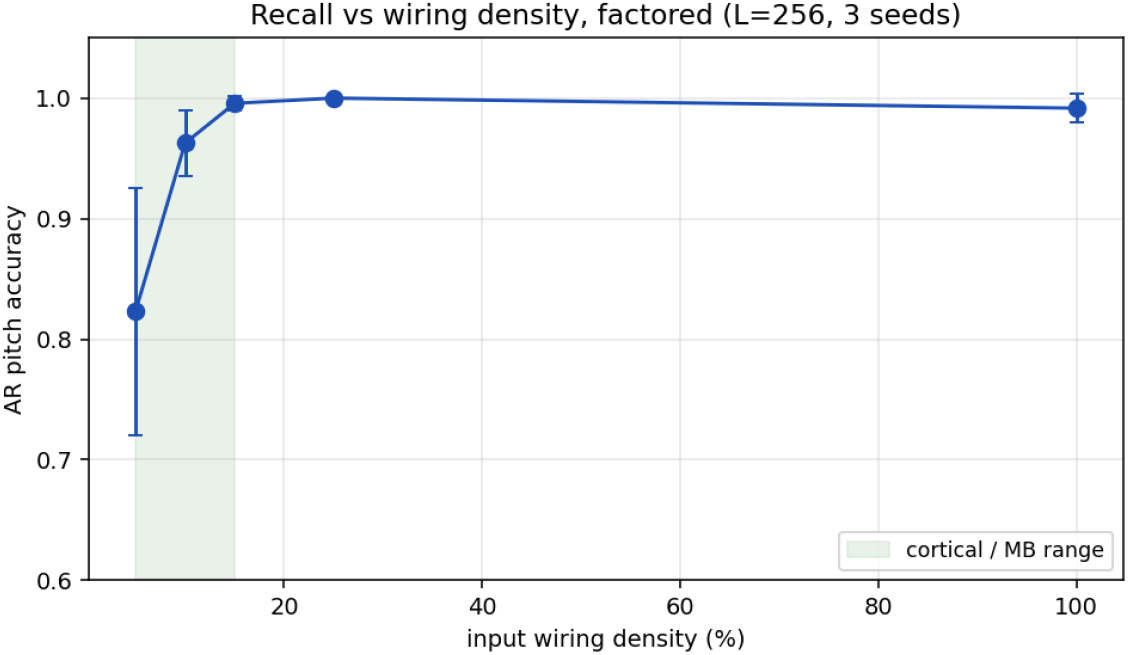
Autoregressive pitch accuracy vs. input-connectivity density at *L* = 256, factored encoding (mean ± std over 3 seeds). Teacher-forced storage is exact from dense to the 10% default; recall is robust across the dense-to-∼10% range (the shaded cortical / mushroom-body range), with the differences among those densities within seed variance, and falls only at 5%. Cutting wiring to cortical density carries no measurable cost.

### 4.6 Structure vs. a random corpus

To separate the contribution of musical *structure* from that of the *memory*, we build 50 random “pieces” matched to the corpus in size and onset/sustain statistics, collision-free by construction, and run the identical sweep at matched (*L, k*), comparing real and random at the same seed and reporting 5 seeds (Table 4). Three findings emerge. First, teacher-forced storage saturates at 1.000 for *both* corpora by *L* = 256: exact storage is a property of the substrate, not of musical structure. Second, final autoregressive fidelity is also structure-independent: by *L* = 512 both reach a perfect 1.000 ± 0.000. Third, wherever the two differ at sub-saturation capacity, musical structure is *not* an advantage: at *L* = 128 the random corpus is recalled better than the real one (0.518 ± 0.039 vs 0.410 ± 0.022), plausibly because diverse random contexts spread load across the sparse code whereas real music’s reused sub-phrases crowd it, and at *L* = 256 the random corpus is again at least as good (0.970 ± 0.032 vs 0.938 ± 0.040, overlapping). We therefore find no evidence that musical structure aids recall on this benchmark: the substrate reaches the same recall fidelity for structured and unstructured data once given enough units, and the only differences, at sub-saturation capacity, slightly favour the unstructured corpus.

**Table 4.** Real vs. a size/onset-matched random corpus at matched capacity and sparsity, factored encoding (120 epochs, mean ± std over 5 seeds, seed-matched between corpora). TF is exact for both above the storage floor.

| Corpus | $L$ | $k$ | TF pitch | AR pitch |
| --- | --- | --- | --- | --- |
| real | 128 | 15 | 0.750 | $0.410 \pm 0.022$ |
| random | 128 | 15 | 0.986 | $0.518 \pm 0.039$ |
| real | 256 | 30 | 1.000 | $0.938 \pm 0.040$ |
| random | 256 | 30 | 1.000 | $0.970 \pm 0.032$ |
| real | 512 | 60 | 1.000 | **$1.000 \pm 0.000$** |
| random | 512 | 60 | 1.000 | **$1.000 \pm 0.000$** |

### 4.7 Capacity to store every melody, including the held-out ones

Having established exact storage of the training set, we asked the complementary question: can the substrate store *every* melody in the dataset, including the 12 held-out ones, if they too were presented for memorization? It can store all but one. The single exception is the spanish song *El cocherito leré*, and the reason is structural rather than a capacity limit: its verse `G G G E` | `F E D D` | `E E E C` recurs within the same melody with two different continuations (`D D C C` on the internal repeats, `F E D C` at the final cadence), so an identical 3-bar context must map to two different next bars. No deterministic next-bar memory can satisfy both demands at once; the best any such model can do is the more frequent continuation, leaving a fixed residual on that one melody. This is the only one-to-many context mapping anywhere in the 62-melody dataset, as verified by an exhaustive scan of all windows. The episode is a small illustration of the boundary between a memory and a function: the limit here is not how much the network can store, but whether the mapping it is asked to store is a function at all.

## 5 Discussion

### 5.1 A minimal recipe that survives Dale’s law

The minimal recipe is a signed local error and a sparse code; what this work adds is that the recipe survives a Dale-compliant substrate. Imposing Dale’s law (signed populations, non-negative weights), explicit quantal synapses, and a sparsely wired frozen encoder does not break the memory; it only raises the latent capacity needed for perfect free-running recall to a modest expansion (*L* = 512 suffices), after which recall is exact. The encoder is still not learned, its wiring can be cut to cortical density, and dense connectivity and a learned front end both fail to help (the latter actively hurts). The working circuit is thus a sparse random expansion read out by a local signed rule under Dale’s law: structurally the fly mushroom-body motif [5, 6]. The biology this requires is modest, no backpropagation, no weight transport, no learned front end, and it is moreover consistent with Dale’s law and multi-contact synapses, which a freely signed read-out would not be.

### 5.2 Is it memorizing or learning? A precise answer

By *memorization* we mean storing specific items for recall; by *learning* (in the generalization sense) we mean acquiring a rule that extends to unseen inputs. On this distinction the model is unambiguously a *structured sparse associative memory*, and three observations pin that down, each holding under Dale’s law.

First, its capacity is governed by sparse-code geometry, as expected for associative memories. The storage floor lies between *L* = 128 and *L* = 256 within an approximately fixed sparse-code regime (*k/L* ≈ 0.12), showing that neither sparsity alone nor raw parameter count explains the transition: storage requires a sufficiently high-dimensional sparse code with enough active units to separate the 293 context-to-next-bar associations [15–17]. This behaviour is consistent with a sparsity-controlled associative memory.

Second, recall is *consistent with attractor-like dynamics* rather than with context indexing. At the working capacity the teacher-forced/autoregressive gap is exactly zero: feeding the model its own outputs back as context does not cause drift, so a stored melody is recovered as a stable trajectory under the model’s own dynamics. A lookup table indexed by the exact 3-bar context would have no reason to be stable under its own (imperfect) outputs; an associative memory that completes patterns toward stored attractors would [18]. The way the gap closes with capacity (Sec. 4.3) is consistent with basins widening until every roll-out stays on the stored manifold, though we do not measure the basin geometry directly.

Third, the code is structured enough to reject novelty. Twelve never-seen melodies are flagged with *zero overlap*, the known-song residual being identically zero while every held-out melody has a residual of at least 0.11: the learned sparse code places unfamiliar inputs outside the stored manifold. A pure per-item table would not by itself provide a graded residual geometry over near and far inputs; the present model does, and that is what an associative memory responding to partial and approximate matches affords [19]. This is the one sense in which the system “generalizes”: it generalizes the *boundary* of what it has stored.

What the model does *not* do is generalize in the predictive sense: it cannot produce or classify a melody it never saw; it can only reproduce stored ones and flag the rest as novel. For a memory device this is not a deficiency: over-parameterization relative to the corpus is appropriate and intended, “overfitting” is the objective, and the figures of merit are storage capacity and recall fidelity, not held-out predictive accuracy.

### 5.3 Relation to overparameterized networks as associative memories

Radhakrishnan, Belkin, and Uhler [20] showed that associative-memory behaviour can emerge in standard overparameterized networks trained with ordinary gradient methods: an autoencoder trained to very low reconstruction error often makes training samples attractors, so iterating the learned map from a corrupted input recovers a stored example. They emphasize that interpolation alone is not sufficient for memory; recoverability requires basins of attraction. The result is particularly relevant for sequences: training a network with *f* (*x*^(*i*)^) = *x*^(*i*+1)^ makes the stored sequence a stable discrete limit cycle. Our setting differs in three ways: the map is a shallow sparse random expansion with a local signed-XOR read-out under Dale’s law, not an overparameterized autoencoder optimized by global gradient descent; recall is an autoregressive context-to-next-bar transition; and the capacity transition is controlled by the geometry of the sparse code, not by *k/L* alone. Nevertheless the phenomena are analogous, and reliable free-running recall corresponds to convergence onto stored sequence trajectories. We thus show that a much more constrained, biologically lawful circuit obtains the same memory behaviour when two interpretable ingredients, signed local error and sparse coding, are imposed explicitly: a constructive, local, sparse, *Dale-compliant* implementation of the overparameterized-memory view. The distinctive message is that recoverability can be achieved without backpropagation, without learning the front end, and without violating Dale’s law, provided a sparse random code is read out by a directional local error.

### 5.4 Relation to biologically plausible fast sequence memories

The literature contains one directly musical precedent and several non-musical but highly relevant fast-memory mechanisms. The closest musical line is the BrainCog sequence-memory model: a spiking architecture with Izhikevich-like neurons, separate content and timing components, dynamically generated excitatory/inhibitory connections, and STDP-modulated synapses, tested on MIDI melodies with goal- and context-based retrieval [21]. A later extension used the same general framework for stylistic composition, coupling a sequential musical-memory subsystem to a more hierarchical knowledge subsystem [22]. These works are important because they show that symbolic musical material can be represented and recalled by local spiking circuits; they are not, however, a drop-in comparison for the present benchmark, because they use a richer spiking architecture, explicit goal/context cues and timing modules, and do not isolate a small 1-, 3-, or 5-presentation regime on a controlled corpus of children’s melodies. Related BrainCog work on arbitrary sequence production further shows that reward-modulated STDP can support the reconstruction of structured symbol sequences, again in a spiking rather than a shallow encoder–decoder setting [23].

The strongest evidence for genuinely rapid storage comes from non-musical memory models. H-Mem places a hetero-associative Hebbian memory inside a trainable network and shows that such a module can store stimulus associations in one shot [24]. Biological key–value memory networks make a similar point using three-factor plasticity rules, extending the key–value idea to auto-association, hetero-association and sequence learning [25]. A spiking variant shows that rapid Hebbian key–value plasticity can enrich SNN computation and support one-shot, cross-modal and language-like memory tasks [26]. Other approaches address sequential recall more directly: temporal predictive coding casts sequence memory as a biologically plausible predictive-coding/asymmetric-Hopfield mechanism [27], while predictive attractor models use sparse distributed representations, lateral inhibition and local Hebbian rules to learn streams online, observing each input only once [28]. Bayesian–Hebbian BCPNN models add an older but still relevant attractor-based route to temporal sequence storage [29, 30].

These precedents sharpen the interpretation of our result. First, they separate two problems that are often conflated: learning a machinery that knows how to address and use a memory, and writing a particular episode into that machinery. Some fast-memory systems are hybrid in this sense: a base network may be optimized globally or meta-trained, but the actual memory trace is written locally and rapidly. Our model takes a different point on this spectrum. There is no pre-trained controller and no global gradient rule; the encoder is frozen, the read-out is updated locally, and the substrate obeys Dale’s law. Conversely, our present experiments use a 120-epoch storage budget and therefore should not be advertised as a one-shot benchmark. What they establish is more specific: a Dale-compliant, sparse, locally updated encoder–decoder can behave as an exact sequence memory on a small melody corpus once capacity is sufficient. The natural next comparison is therefore not a larger RNN baseline, but a strict exposure-limited protocol – one, three, five and ten presentations per melody – against STDP music-memory, H-Mem/key–value, temporal-predictive coding and PAM-style attractor baselines.

### 5.5 Parameters, information content, and the cost of Dale’s law

The corpus’s 50 melodies comprise 1,772 beat slots (1,445 onsets); their content is about 6,100 bits by gzip (and ∼ 8,100 bits under a uniform pitch×attack bound). At the storage floor *L* = 256 the decoder credit-assigns ∼ 14,300 connection bundles and the 10% encoder has only ∼ 4,300 fixed random synapses; the learned budget is thus within a small factor of the corpus’s own information content, as expected of an associative memory rather than a function approximator fit far past the data. Two consequences follow. The binding constraint is representational, not informational: recall collapses below the *L* ≈ 256 threshold despite ample bit-capacity at every *L*, so the threshold is the geometry of the sparse code, not the bit budget. And the extra capacity Dale’s law demands (the headline model uses *L* = 512, ∼ 28,700 decoder bundles) is the price of a positive-only read-out: it does not change *what* is stored, only how many units are needed to route suppression cleanly enough for a perfect roll-out.

### 5.6 Capacity scaling, reconsidered

The model’s autoregressive rise from 0.27 to 1.0 as *L* grows is, against the teacher-forced storage that is exact from *L* = 256, the picture of a positive-only read-out slowly buying back the margin a freely signed decoder would have for free, until at *L* = 512 it matches it exactly. Capacity curves are uninterpretable without the minimal-storage baseline that says how much capacity the task actually needs: here that baseline is the exact teacher-forced storage, and the autoregressive curve measures the overhead of Dale’s law specifically.

### 5.7 The teacher-forced/autoregressive gap

The TF–AR gap is a useful single-number signature of clean storage: at the working capacity (*L* = 512) it is exactly zero, so free-running recall does not drift. Just above the storage floor the gap is small but nonzero (1.000 vs 0.963 at *L* = 256) and closes to zero by *L* = 512; below the floor, where teacher forcing itself is sub-exact, free-running recall falls much further (AR 0.410 at *L* = 128). The gap therefore tracks how cleanly a configuration has stored the corpus given its capacity, not a property of any one architecture.

### 5.8 Limitations

The corpus is small and discrete by design, chosen only as a controllable example of sequence memory; conclusions about generalization, larger or noisier data, or long-term retention are out of scope, and the children’s-song framing is incidental. Recognition and teacher-forced storage saturate, so autoregressive recall and known–novel separation carry the discriminative signal. This version reports the local Dale’s-law model only; a matched non-biological reference (a BPTT-trained LSTM with scheduled sampling, the natural exposure-bias-corrected baseline) on this exact corpus and substrate is left for a companion measurement, and we make no head-to-head claim here. The novelty separation is reported as zero overlap rather than in units of the known-song standard deviation because, at exact storage, the known-song residual variance is zero and the standardized separation is degenerate. Finally, the model is a rate-coded, discrete-time abstraction; a spiking realization of the signed-XOR motif is future work.

### 5.9 Future work: a more brain-like pitch code

A clear next step concerns the input/output representation. Pitch is currently encoded localistically, as a one-hot over the twelve chromatic classes, so the twelve pitches are mutually orthogonal and equidistant and the representation discards the metric structure of pitch: neighbouring-semitone similarity, octave-equivalent chroma, and the interval relations on which human melody recognition primarily rests. The brain does not represent pitch this way: auditory cortex is tonotopically organized with broadly, overlappingly tuned neurons [31], and pitch perception carries a circular chroma component [32]. Several more biologically grounded codes, all compatible with the local signed rule provided the output stays binary, are natural to explore: a tonotopic population code in which each pitch is a contiguous active arc so that neighbours share units and the code reflects pitch distance (placed on a circle to capture octave equivalence); and a relative, interval-based code conferring transposition invariance, consistent with the contour- and interval-based memory humans use for melodies [33]. A one-hot code is in fact near-optimal for pure memorization, precisely because it makes stored patterns maximally separable; a similarity-preserving code trades some separability for metric structure, and that trade is exactly the lever that could move the system from storing specific melodies toward genuine generalization, robustness to single-semitone errors and recognition of transposed tunes. Testing one-hot against population and relative encodings, with an explicit transposition probe, is therefore a direct way to ask whether a sparse associative memory equipped with a structured input code begins to *learn* rather than only memorize. We leave this to future work.

## 6 Conclusion

Held to a substrate that obeys Dale’s law, uses explicit multi-contact synapses, and reads from a frozen encoder wired at cortical density, a biologically plausible sequence memorizer still needs very little: a signed local error and a sparse code, with the Dale population structure, memorize a small melody corpus and, given enough latent units, recall every tune perfectly under free running, recognizing all 50 and rejecting all 12 novel ones with zero overlap, without backpropagation. The encoder need not be learned and its wiring can be cut to the ∼ 10% density of cortex, with dense wiring no better and a learned front end worse, giving a mushroom-body-like associative memory. Dale’s law does not break the memory; it only sets a higher capacity for perfect free-running recall, the price a positive-only read-out pays to route suppression, which is fully recovered with scale. And the system is best understood not as learning to generalize but as a structured sparse associative memory, one whose capacity is set by sparse-code geometry, whose recall is consistent with attractor-like dynamics, and whose code is just structured enough to know what it has not seen.

## 7 Methods

### 7.1 Encoding

Melodies are parsed into 4*/*4 bars of four beats; each beat carries a pitch class (one of twelve) and an attack state (onset vs. sustain). The next bar is predicted from the previous *K* = 3 bars, encoded as a 168-dimensional vector (12 × slots 14 dimensions: a 12-way one-hot pitch and a separate 2-way one-hot {onset, sustain}). The decoder emits the same 14-dimensional code per slot, so output and input share a single neural format. The 50 training melodies contain no two contexts that map to different next bars; the single melody carrying a one-to-many context mapping is held out (Sec. 4.7).

### 7.2 Model and learning rule

The encoder **W**_enc_ ∈ ℝ^*L×*168^ is a fixed, sparsely wired random projection (∼10% density by default) producing **h**; *k*-WTA retains the top *k* units (*k/L* ≈ 0.12 at the storage floor). Latent units are excitatory or inhibitory (80*/*20); all weights are non-negative and a unit’s output sign is that of its population (Dale’s law). Each decoder connection is an explicit bundle of up to ten quantal contacts; per-beat decoders emit a 14-dimensional code read out by two winner-take-all heads – a 12-way head for pitch and a 2-way head for {onset, sustain} – and are updated by the signed-XOR rule (Eq. 1) applied independently to each head, potentiation filling contacts and LTD pruning the weakest labile ones, with consolidation locking repeatedly potentiated contacts. The encoder is frozen by default; the optional learned variant receives the output error through a fixed random matrix **B** (feedback alignment), with the sparse-wiring mask reapplied after each update so absent connections never re-grow. No global loss or backpropagation is used.

### 7.3 Conditions and budget

All conditions share one code base and differ only in switch settings (Dale on/off; signed/ unsigned; *k*-WTA on/off; multi-synapse on/off; consolidation, LTD, divisive inhibition, recognition head, encoder learning; connectivity density). Capacity, connectivity, and ablation sweeps use a 120-epoch budget (conservative; metrics plateau by ∼60 epochs, Sec. 4.4), the ablation at *L* = 256, *k* = 30, at a fixed learning rate with no early stopping.

### 7.4 Evaluation

TF accuracy predicts each bar from the true context; AR accuracy feeds the model’s own predictions back. Recognition votes the per-window classifier head against a per-model threshold. Novelty is the residual teacher-forced prediction error; a tune is flagged novel when its residual exceeds the known-song mean by 2*σ*. At exact storage the known-song residual is zero, so novelty separation is reported as overlap (zero) rather than in standard-deviation units. AR is reported as mean ± std over three seeds where noted; TF and recognition are deterministic at saturation.

## Data and code availability

The corpus encoding, the Dale’s-law model (including the sparse-connectivity mask, the quantal multi-synapse decoder, and the mechanism switches), and the drivers that regenerate every table and figure (the capacity, ablation, wiring, and random-corpus sweeps) are available at https://github.com/jesusmarcodelucas/musicmem-engrammer. The signed-XOR substrate this work builds on is described in publicly available work [1, 2], with reference code at https://github.com/MariaPFdez/XOR_classic.

## Acknowledgments

This work was supported by the ENGRAMMER project (IASOMM2024007), funded by the EU NextGeneration programme under the Recovery and Resilience Facility (RRF).

## A Encoding comparison: factoring pitch and attack

The main text uses the *factored* encoding: each beat is two clean one-hot populations, a 12-way pitch code and a separate 2-way {onset, sustain} code (14 dimensions), and the decoder emits the same 14-dimensional code through two winner-take-all heads. We compare it against two alternatives that encode the identical information differently: *attack* (13 dimensions), which glues a single onset *bit* onto the pitch one-hot, and *dual* (24 dimensions), which tags onset within a pitch-specific block (sustained[12] + onset[12]). Table 5 reports autoregressive pitch accuracy for all three on the real corpus and on the matched random corpus, over 5 seeds.

**Table 5.** Autoregressive pitch accuracy by input encoding, latent size, and corpus (mean ± std over 5 seeds; *k/L* ≈ 0.12). Teacher-forced storage is exact for all three encodings by *L* = 256. The factored encoding reaches perfect recall at *L* = 512.

| Encoding | $L$ | AR (real) | AR (random) |
| --- | --- | --- | --- |
| attack (13-d) | 128 | $0.369 \pm 0.019$ | $0.501 \pm 0.021$ |
| dual (24-d) | 128 | $0.495 \pm 0.038$ | $0.518 \pm 0.040$ |
| factored (14-d) | 128 | $0.410 \pm 0.022$ | $0.518 \pm 0.039$ |
| attack (13-d) | 256 | $0.677 \pm 0.017$ | $0.629 \pm 0.015$ |
| dual (24-d) | 256 | $0.740 \pm 0.032$ | $0.699 \pm 0.015$ |
| factored (14-d) | 256 | **$0.938 \pm 0.040$** | **$0.970 \pm 0.032$** |
| attack (13-d) | 512 | $0.888 \pm 0.049$ | $0.892 \pm 0.027$ |
| dual (24-d) | 512 | $0.991 \pm 0.008$ | $0.953 \pm 0.025$ |
| factored (14-d) | 512 | **$1.000 \pm 0.000$** | **$1.000 \pm 0.000$** |

Three points stand out. First, the factored encoding is decisively the best above the storage floor: it reaches a perfect 1.000 autoregressive recall at *L* = 512 on both corpora – the capacity at which the attack code is still at 0.888 and the dual code at 0.991 – and it already attains ∼ 0.95 at *L* = 256 where attack and dual sit near 0.68–0.74. It thus roughly halves the latent capacity needed for flawless free-running recall. Second, this gain is not bought with parameters: the factored decoder is 4 × 14 × *L* learned bundles, only one output unit per beat more than attack’s 4 × 13 × *L*, whereas a format-closed dual decoder would need 4 × 24 × *L*. Third, the advantage is representational, not musical: the factored code also recalls the *random* corpus better, so the benefit comes from disentangling pitch and rhythm into two symmetric one-hots (every slot has exactly two active units, with sustain a positive state) rather than from any property of the melodies. The attack code is asymmetric (onset slots fire two input units, sustain slots one, and “sustain” is the absence of a bit) and the dual code, while richer, is not format-closed unless its decoder is doubled; the factored code is symmetric, format-closed, and the most biologically natural, separating pitch from timing into distinct populations.

